# Accelerated epigenetic aging and inflammatory/immunological profile (ipAGE) in patients with chronic kidney disease

**DOI:** 10.1101/2021.07.23.453588

**Authors:** Igor Yusipov, Elena Kondakova, Alena Kalyakulina, Mikhail Krivonosov, Nadezhda Lobanova, Maria Giulia Bacalini, Claudio Franceschi, Maria Vedunova, Mikhail Ivanchenko

**Affiliations:** Institute of Information Technologies, Mathematics and Mechanics, Lobachevsky State University, Nizhny Novgorod, Russia; Institute of Biology and Biomedicine, Lobachevsky State University, Nizhny Novgorod, Russia; IRCCS Istituto delle Scienze Neurologiche di Bologna, Bologna, Italy

**Keywords:** Chronic kidney disease, Accelerated aging, DNA methylation, Inflammation

## Abstract

Chronic kidney disease (CKD) is defined by reduced estimated glomerular filtration rate (eGFR). This failure can be related to a phenotype of accelerated aging. In this work we considered 76 subjects with end-stage renal disease (ESRD) and 83 healthy controls. We evaluated two measures that can be informative of the rate of aging, i.e. whole blood DNA methylation using the Illumina Infinium EPIC array and plasma levels of a selection of inflammatory/immunological proteins using Multiplex Immunoassays. We demonstrated accelerated aging in terms of the most common epigenetic age estimators in CKD patients. We developed a new predictor of age based on inflammatory/immunological profile (ipAGE) and confirmed age acceleration in CKD patients. Finally, we evaluated the relationship between epigenetic age predictors and ipAGE and further identified the inflammatory/immunological biomarkers differentially expressed between cases and controls. In summary, our data show an accelerated aging phenotype in CKD patients sustained by inflammatory processes.

## Introduction

The nature of aging and its key mechanisms are still a matter of debate^1^. However, there is no doubt that many pathologies, both spontaneously occurring and genetically determined, are age-related, to name cancer, endocrine and cardiovascular diseases, growth and functional disorders of bones and joints.

Aging and age-related diseases share some basic mechanistic pillars that largely converge on inflammation. This has been conceptualized as inflammaging, the long-term result of the chronic physiological stimulation of the innate immune system, which can become damaging during aging^2,3^. Inflammaging poses a highly significant risk factor for both morbidity and mortality in the elderly people, as most if not all age-related diseases share an inflammatory pathogenesis^4,5^.

Recently, Alpert, Davis, Shen-Orr and colleagues have demonstrated the comprehensive remodeling of the immune cell composition with age, and developed an aggregated score IMM-AGE, associated with both chronological and DNA methylation age, as well as cardiovascular disease risk^6^. Similarly, Sayed et al. recently proposed an inflammatory aging clock (iAge), based on circulating inflammatory proteins encompassing cytokines, chemokines and growth factors, that has been shown to be informative of multimorbidity, immunoscenescence and cardiovascular aging^7^.

The inflammatory clocks are part of a larger family of biomarkers of aging^8^, in which epigenetic clocks have raised to the fame^9^. In general, epigenetic clocks have predictive power of chronological age higher than other biomarkers, including immunological ones. Numerous studies showed that the aging process and the development of age-related syndromes and diseases and syndromes are accompanied by epigenetic changes in DNA^10–18^, and may depend on external factors such as alcohol, smoking, microbiome profile, region of residence, and diet^19–21^. Each of these factors contributes differently to epigenetic changes and age acceleration.

A particularly important risk brought by senescence and aging of the immune system is the Chronic kidney disease (CKD), a highly prevalent (10-13% of the population) and irreversible pathology^22^. Progressing to the end-stage renal disease (ESRD), it is considered as a major issue for public health^23,24^. Between hemodialysis sessions, the organism gets into the state of permanent intoxication, because the patients’ kidneys have completely lost their detoxification function. Intoxication emerging from impaired renal function can activate the processes of chronic inflammation. Similarly, to the development of oxidative stress and atherosclerosis, the self-proteins get modified by toxic metabolic products, lose their functional role, and become considered by the innate immune system as antigens, further stimulating the processes of inflammaging.

So far, some studies have reported accelerated aging phenotypes in ESRD^25–28^. A recent study demonstrated the association between epigenetic clocks and kidney disease, but it did not include ESRD patients^29^.

In the present study we investigated for the first time epigenetic and immunological/inflammatory biomarkers of age in ESRD patients.

## Methods

### Participants

All recruited individuals provided written, informed consent, and this study was approved by the local ethics committee of Nizhny Novgorod State University. All participants were recruited from Nizhny Novgorod, Russia. The dataset has an almost equal distribution by sex: 81 women and 78 men (Figure 1(a)). The dataset includes 76 subjects with 5-stage CKD (ESRD) and 83 controls (Figure 1(b)). The age range of subjects ranges from 24 to 89 years. Figure 1(c) represents the ratio between controls and cases, males and females. Some of the subjects in the dataset (mainly ESRD patients) have comorbidities. Figure 1(e) shows the most representative diseases in the group of subjects with ESRD, coded according to the International Classification of Diseases (ICD-10)^30^. In addition to stage 5 Chronic kidney disease (N18.5), which occurs in all patients in the respective group, the most common diseases are anemia (D63.8) in 69 subjects, hypertension (I15.8) in 65 subjects, secondary hyperparathyroidism (E21.1) in 48 subjects. Figure 1(f) shows the most representative medicines (codes in Anatomical Therapeutic Chemical Classification System^31^), which consume patients with ESRD. Most of them take folic acid (B03BB01) and cyanocobalamin (B03BA01). All biological data from the subjects with the disease were taken within 30 minutes after the dialysis procedure. Supplementary Table 1 contains full information about the Russian population dataset (sheet “Main” provides information about age, sex, group, time on dialysis, sheet “Diseases” contains information about diseases, sheet “Drugs” contains information about drugs taken).

**Figure 1.**
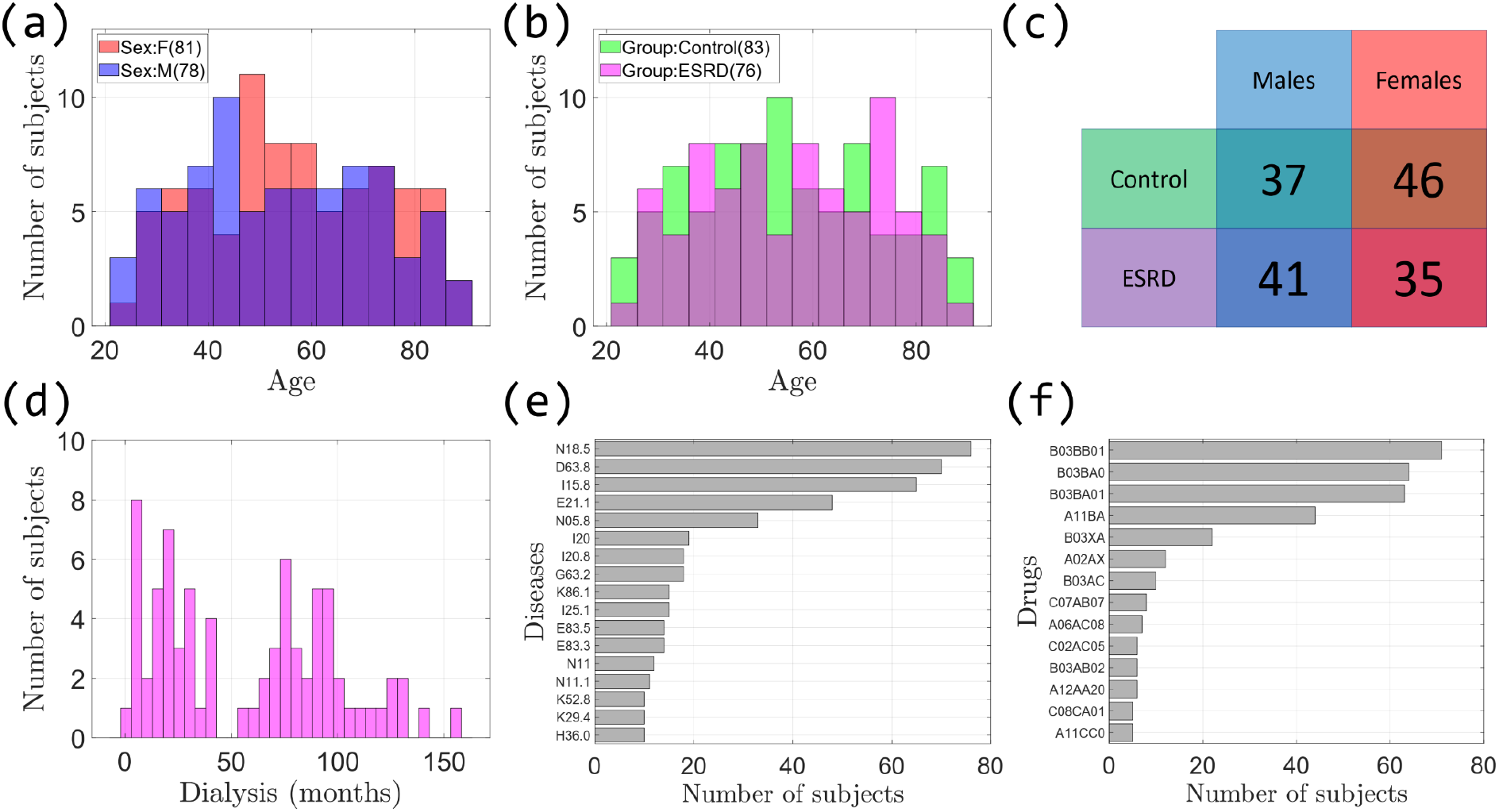
Russian population dataset characteristics. (a) Distribution of men and women by age; (b) Distribution of controls and subjects with the ESRD by age; (c) Summary table by group and sex; (d) Distribution subjects with the disease by time on dialysis, measured in months; (e) The most representative diseases; (f) The most representative drugs.

### DNA methylation quantification and quality control

Phenol Chloroform DNA extraction was used. DNA was quantified using the DNA Quantitation Kit Qubit dsDNA BR Assay (Thermo Fisher Scientific) and 250 ng were bisulfite-treated using the EpiMark Bisulfite Conversion Kit (NEB) with case and control samples randomly distributed across arrays. The Illumina Infinium MethylationEPIC BeadChip^32^ was used according to the manufacturer’s instructions. This platform allows measuring DNA methylation levels from a total number of 866,836 genomic sites, with single-nucleotide resolution. DNA methylation is expressed as β values, ranging from 0 for unmethylated to 1 representing complete methylation for each probe. Raw data were pre-processed as follows. First, probes with a detection p-value above 0.01 in at least 10% of samples were removed from the analysis. Second, probes with a beadcount less than three in at least 5% of samples, were removed from the analysis. Third, all non-CpG probes were excluded from the results^33,34^. Fourth, SNP-related probes were removed from the analysis^34^. Fifth, multi-hit probes were removed^35^. Sixth, all probes located in chromosome X and Y were filtered out. As a result, 733,923 probes remained for the analysis. All samples have less than 10% of probes with a detection p-value above 0.01. Functional normalization of raw methylation data was performed using the minfi^36^ R package.

### Multiplex Assay Kits

The analysis was performed on plasma using the K3-EDTA anticoagulant, without hemolysis and lipemia. Plasma from matched subjects was thawed, spun (3000 rpm, 10 min) to remove debris, and 25 μl collected in duplicate. Plasma with antibody-immobilized beads was incubated with agitation on a plate shaker overnight (16-18 hours) at 2-8 °C. The Luminex® assay^37^ was run according to the manufacturer’s instructions, using a custom human cytokine 46-plex panel (EMD Millipore Corporation, HCYTA-60K-PX48). The panel included: sCD40L (CD40LG), EGF, Eotaxin (CCL11), FGF-2, FLT-3L, Fractalkine (CX3CL1), M-CSF (CSF1), GROα (CXCL1), IFNα2, IFNγ (IFNG), IL-1α, IL-1β, IL-1RA, IL-2, IL-3, IL-4, IL-5, IL-6, IL-7, IL-8, IL-9, IL-10, IL-12 (p40), IL-12 (p70) (IL12Bp70), IL-13, IL-15, IL-17A, IL-17E/IL-25 (IL25), IL-17F, IL-18, IL-22, IL-27, IP-10 (CXCL10), MCP-1 (CCL2), MCP-3 (CCL7), M-CSF, MDC (CCL22), MIG (CXCL9), MIP-1α (CCL3), MIP-1β (CCL4), PDGF-AA, PDGF-AB/BB (PDGFB), TGFα, TNFα, TNFβ (LTA), VEGF-A. Assay plates were measured using a Magpix (Milliplex MAP). Data acquisition and analysis were done using a standard set of programs MAGPIX®. Data quality was examined based on the following criteria: The standard curve for each analyte has a 5P R2 value > 0.95. To pass assay technical quality control, the results for two controls in the kit needed to be within the 95% of CI (confidence interval) provided by the vendor for >40 of the tested analytes. No further tests were done on samples with results out of range low (<OOR). Samples with results that were out of range high (>OOR) or greater than the standard curve maximum value (SC max) were not tested at higher dilutions without further request. Supplementary Table 2 contains Multiplex immunohistochemistry biomarkers data.

### Biochemical markers

Two types of anticoagulants (K3-EDTA and Li-heparin) were used. To identify possible reasons for the difference between biological and chronological ages, the biological age estimation using the Levine’s model^38^ was carried out in all the samples obtained. For this purpose, a CBC was performed using the Abacus Junior 30 semi-automatic analyzer and a set of biochemical indicators using the analyzer StatFax 3300 and diagnostic kits DIAKON-DS (Russia). Supplementary Table 3 stores Phenotypic Age and the blood biochemistry markers which were used for calculating Phenotypic Age^38^.

To investigate the effect of dialysis on the values changing of blood biochemistry parameters, which were used to calculate the Phenotypic Age, we considered data for 5 people who are not included in the main dataset (Supplementary Figure 1, Supplementary Table 4). Creatinine shows the highest change after dialysis.

### Age acceleration

The work considered four types of epigenetic age: DNAmAgeHannum^39^, DNAmAge^40^, DNAmPhenoAge^38^, DNAGrimAge^41^. The DNAmAgeHannum quantitative aging model measures the aging rate of human methylome in whole blood. The DNAmAge multi-tissue age predictor allows estimating the age of DNA methylation in various tissues and cells. The biomarker of aging DNAmPhenoAge was developed considering composite clinical measures of phenotypic age^38^, while the DNAmGrimAge is a composite biomarker based on DNAm surrogates of seven plasma proteins and of smoking history. Data for all 4 models were obtained using Horvath’s calculator^42^ and presented in Supplementary Table 5.

To calculate the acceleration of the epigenetic age of the subjects considered in the study, first, for each type of epigenetic age, a linear regression model which estimates chronological age was built only on the group of control subjects. Epigenetic age acceleration is a residual from this linear model. All models were built using statsmodels^43^ package in Python programming language. Mann–Whitney U test from the scipy^44^ Python package was applied to analyze the statistically significant difference of epigenetic age acceleration between controls and subjects with the disease. The same approach was used to analyze the age-acceleration of Phenotypic Age, based on nine biomarkers and chronological age^38^.

We also calculated a measure of intrinsic epigenetic age acceleration (IEAA), which characterizes “pure” epigenetic aging effects without the influence of differences in blood cell counts^15,16^. IEAA is the residual resulting from a multivariate regression model of DNAm age on chronological age and blood immune cell counts (naïve CD8+ T cells, exhausted CD8+ T cells, plasma B cells, CD4+ T cells, natural killer cells, monocytes, and granulocytes). The model was built only on the group of control subjects using statsmodels^43^ package in Python programming language. We have calculated a measure of extrinsic epigenetic age acceleration (EEAA), which characterizes epigenetic aging in immune-related components^15,16^. EEAA is the residual resulting from a univariate model that regressed the weighted average of the epigenetic age measure from Hannum and three estimated measures of blood cells (naïve (CD45RA+CCR7+) cytotoxic T cells, exhausted (CD28-CD45RA-) cytotoxic T cells, plasma B cells) on chronological age. The model was built only on the group of control subjects using statsmodels package in Python programming language. The distributions of blood cell counts, obtained from Horvath’s calculator^42^, are in Supplementary Figure 2.

### Immunology clock

Using immunology Multiplex assays, we applied the Elastic Net regression model (sklearn^45^ package in Python) where chronological age was regressed on 46 Multiplex biomarkers to select variables for inclusion in our immunology age score (ipAGE). K-fold cross-validation (K=5) was employed to select the optimal parameter alpha, which multiplies the penalty terms (lambda parameter in glmnet^46^ R package). The searching process for the optimal value of this parameter was performed on an equidistant logarithmic grid: {10^−5^, …, 10^−2^, …, 10^1^}. Parameter l1_ratio, which is a mixing of L1 and L2 regularization and corresponds to alpha in the glmnet R package was taken as 0.5. Thus, for each set of parameters, the entire dataset was divided 5 times into training (80%) and test (20%) sets (sliding window of the test set). To calculate the ipAGEacceleration for each model we firstly build a linear regression which estimates chronological age only on controls. After that, acceleration is calculated as a residual from this model. Mann–Whitney U test was applied to check the difference in age acceleration of ipAGEbetween the groups of subjects.

To compare the results of ipAGEclock, we compare their results from other phenotypic and epigenetic clocks by calculating the Pearson correlation coefficient between corresponding ages, as well as between accelerations relative to chronological age.

For a deeper analysis of age acceleration, we used the Principal Component Analysis (PCA) from sklearn^45^ Python package. This is a linear dimensionality reduction technique that allows us to extract lower-dimensional results from dimensions of different types of age accelerations.

To detect patterns of age acceleration in control subjects and subjects with ESRD, we applied the hierarchical clustering method to construct a dendrogram, where each subject is characterized by a corresponding set of age accelerations. For this, we used the sklearn^45^ Python package.

### Associations of immunology biomarkers

To find the most disease-associated immunology biomarkers, the Mann–Whitney U test (scipy^44^ Python package) was applied to the values of the biomarker, divided into groups (controls vs ESRD). To find biomarkers associated with chronological, phenotypic, epigenetic and immuno ages, the Pearson correlation test was performed (scipy^44^ Python package). As a criterion for a statistically significant relationship between biomarkers and the selected age types, the Pearson correlation p-value of the corresponding biomarkers is used, the values of which are less than 0.05 indicate the presence of a relationship. The same approach was used to find associations between biomarkers and different types of age acceleration, which were defined as residuals from the chronological age.

## Results

### Epigenetic age measures in ESRD patients

To identify the relationship between ESRD and aging processes, the 5 most common epigenetic clocks DNAmAgeHannum^39^, IEAA^15,16^, EEAA^15,16^, DNAmAge^40^, DNAmPhenoAge^38^, DNAGrimAge^41^ were analyzed (Figure 2). Our results demonstrate that three out of four epigenetic clocks (DNAmAge^40^, DNAmPhenoAge^38^, DNAGrimAge^41^) show statistically significant (Mann–Whitney U test p-value below 0.05) age-acceleration in the group of subjects with ESRD (Figure 2(b),(e),(f)), as well as IEAA^15,16^ (Figure 2(c)), with DNAmAgeHannum^39^ and EEAA^15,16^ as exceptions (Figure 2(a),(d)). On average, epigenetic age acceleration in the ESRD group was about 3 years. DNAmAgeHannum showed the smallest increase of 0.6 years (Q1=-1.9, Q3=3.5, *p* = 1. 08*e* − 1); the highest age-related acceleration of 7.0 years (Q1=1.7, Q3=11.2, *p* = 4. 11*e* − 7) was recorded for DNAmPhenoAge.

**Figure 2.**
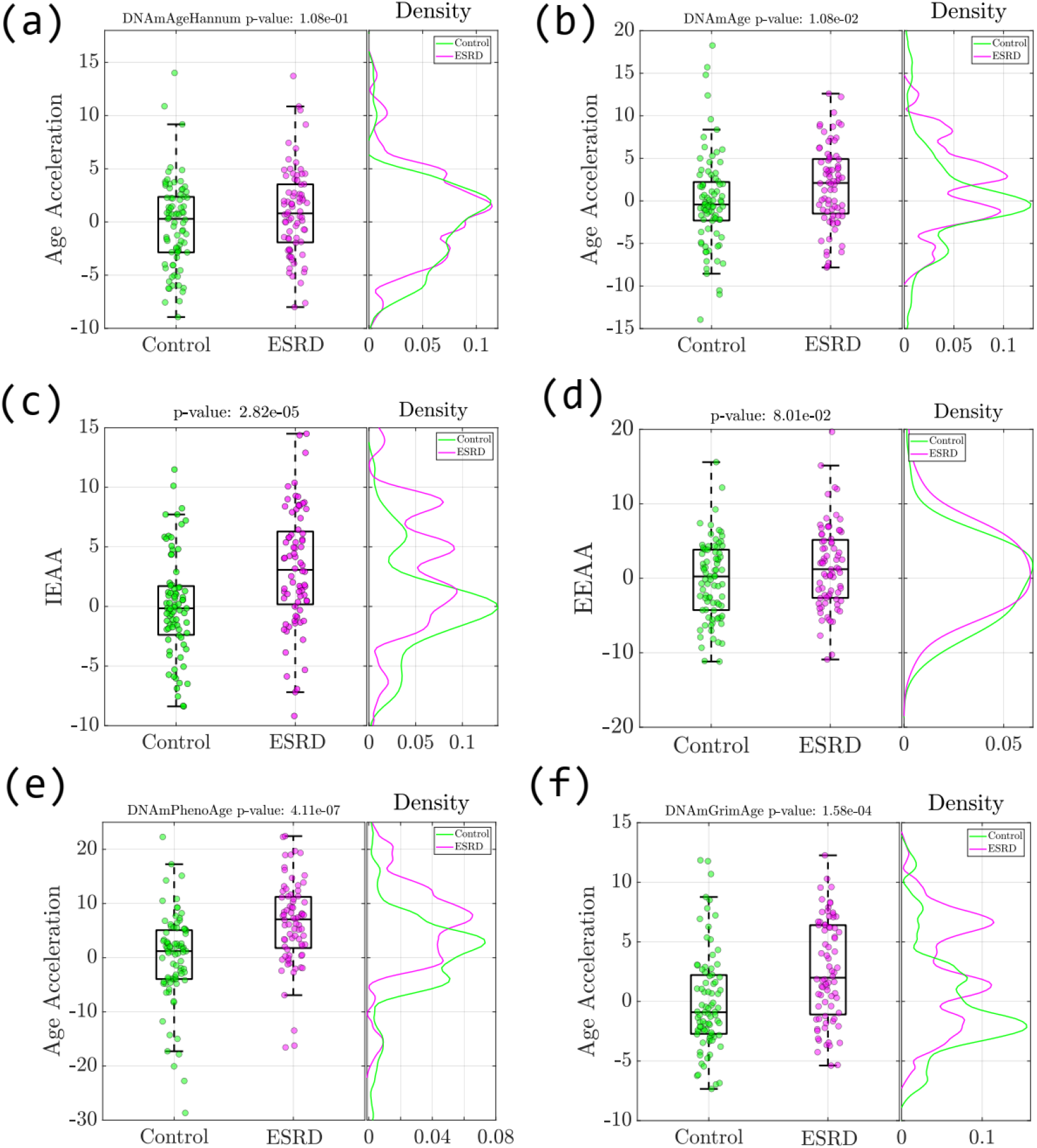
Age acceleration differences in controls and subjects with the ESRD. (a) DNAmAgeHannum^39^; (b) DNAmAge^40^; (c) IEAA^15,16^; (d) EEAA^15,16^; (e) DNAmPhenoAge^38^; (f) DNAmGrimAge^41^.

Next, we considered Phenotypic Age^38^, a biological age estimator based on biochemical and hematological blood parameters, which could be expected to be more sensitive to the pathological changes in the organism. On the one hand, the employed markers are more responsive to various cues than epigenetic ones, not necessarily related to the pathology; on the other hand, it potentially allows to catch earlier the deficiencies of the body homeostatic regulation caused by the pathology. Besides, Phenotypic Age includes biomarkers, which are potentially associated with the disease (for example, creatinine). In result, the Phenotypic age obtained from blood biomarkers proved to manifest age acceleration in patients with ESRD substantially more pronounced than epigenetic age estimators (Figure 3(a)). To get a deeper insight we plot the distributions of biomarkers involved in these clocks for both the control group and patients with ESRD in Figure 3(b). It follows that almost all biomarkers in the ESRD group display significantly different values from the control group.

**Figure 3.**
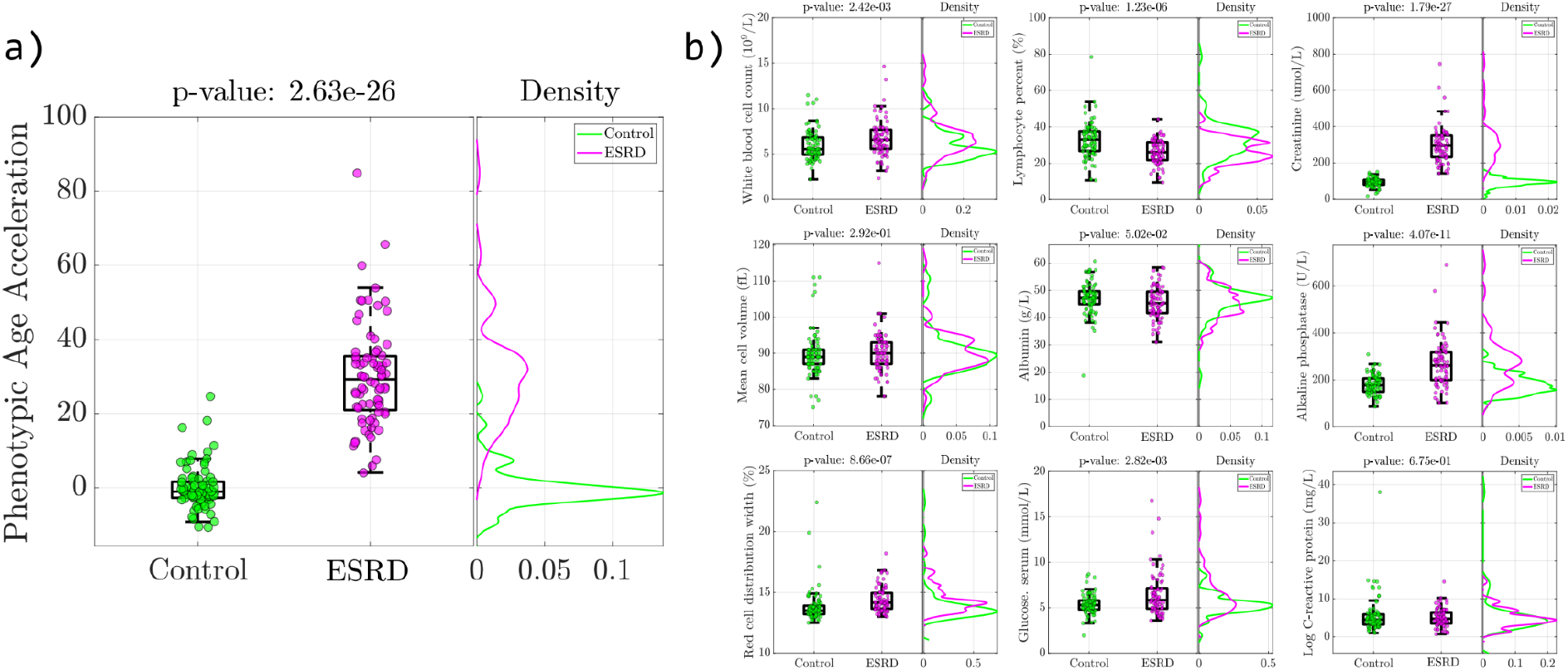
(a) Age acceleration differences in controls and subjects with the ESRD for Phenotypic Age^38^; (b) Distribution of components of Phenotypic Age estimator in controls and subjects with the ESRD.

### Inflammatory/Immunological profile clock (ipAGE)

We evaluated 46 inflammatory/immunological markers in plasma from the subjects involved in the study. Applying Elastic Net regression model in the group of healthy controls (see Methods), we built a model that estimates age based on these biomarkers, termed inflammatory/immunological profile age (ipAGE). The model included 38 out of 46 biomarkers and yielded a determination coefficient 0.79, Mean Absolute Error (MAE) 6.82 years, Root Mean Squared Error (RMSE) 8.17 years in the control group. Supplementary Table 6 reports the list of the proteins included in the ipAGE clock, their model coefficients and the comparison with the iAge from Sayed et al. In particular, 27 plasma proteins are shared between the two predictors. ipAGE values for the considered cohort are listed in Supplementary Table 7. Figure 4(a) illustrates the resulting fit in the control group.

**Figure 4.**
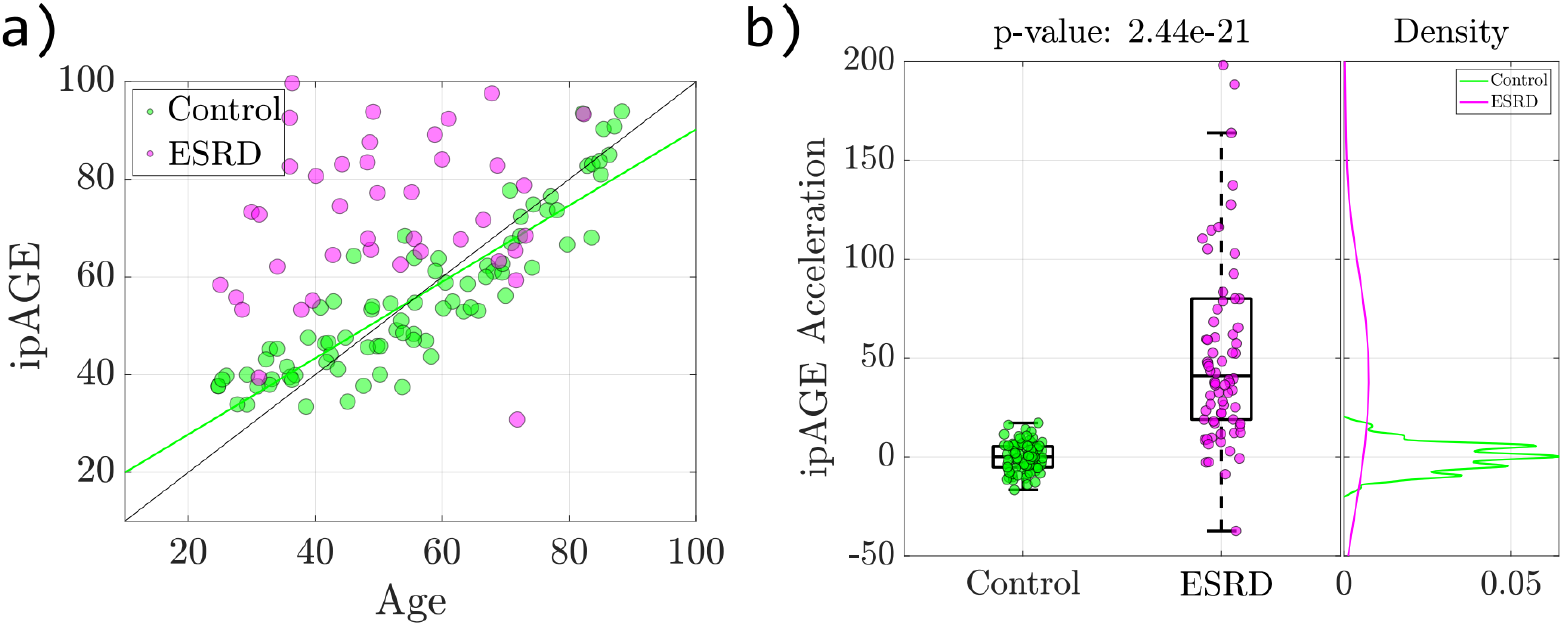
(a) Association between ipAGE and chronological age; (b) ipAGE acceleration in controls and subjects with the ESRD.

As for the epigenetic clocks, we calculated age acceleration as the residuals of the linear regression between ipAGE andand chronological age. Importantly, we found a statistically significant age acceleration in the ESRD group compared to controls, as illustrated in Figure 4(b) (p=2.44e-21). The group of subjects with ESRD exhibited also a large ipAGE variance.

We then investigated the relationship between epigenetic clocks, phenotypic clock and ipAGE. Expectedly, all the clocks showed high correlation with chronological age in control subjects (Figure 5(a)) and between each other. The epigenetic age estimators maintained high correlation with chronological age and among each other also in the ESDR patients, while in this group the ipAGE was weakly correlated with chronological age and the other clocks. This is a consequence of the sensitive response of ipAGE to ESRD with respect to controls and significant intra-group differences between ESRD patients. (Figure 5(b)). Instructively, there was no significant correlation between age accelerations, except correlations between IEAA∼DNAmAgeAcc and EEAA∼DNAmAgeHannumAcc which is due to calculation of IEAA and EEAA being based on DNAmAge and DNAmAgeHannum correspondingly (Figure 5(c-d)). The effect was observed both for the control and ESRD groups, indicating that the same subject can have a positive age acceleration according to some clocks and negative by others.

**Figure 5.**
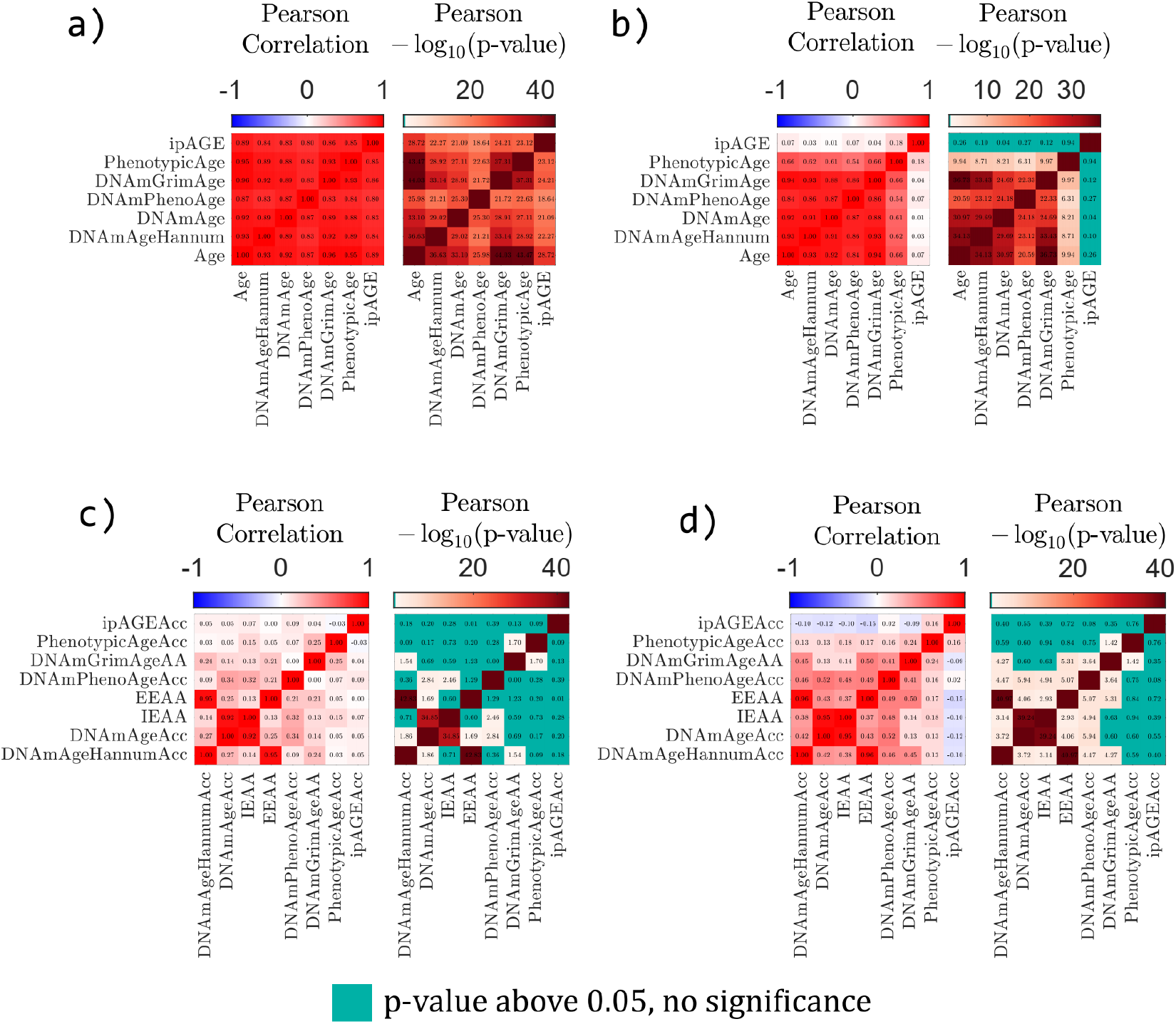
Pearson correlation coefficient between different types of ages and corresponding p-values for (a) control group and (b) ESRD group. Pearson correlation coefficient between different types of age acceleration for (c) control group and (d) ESRD group.

### Association of immunology biomarkers with ESRD status

As a next step, we compared the 46 inflammatory/immunological markers between ESRD and controls Eighteen out of 46 biomarkers (Supplementary Table 8) had a statistically significant association with ESRD (Mann–Whitney U test p-value below 0.05). KEGG analysis indicated that the significantly different immunology biomarkers tended to be enriched in the Toll-like receptor signaling pathway (hsa04620; enrichment ratio: 2.2; nominal p-value<0.01). Figure 6 illustrates the resulting p-values, manifesting the strongest association with disease of CSF1, CXCL9 and IL12Bp40, and exemplifies distributions of the first two biomarkers in control and ESRD groups.

**Figure 6.**
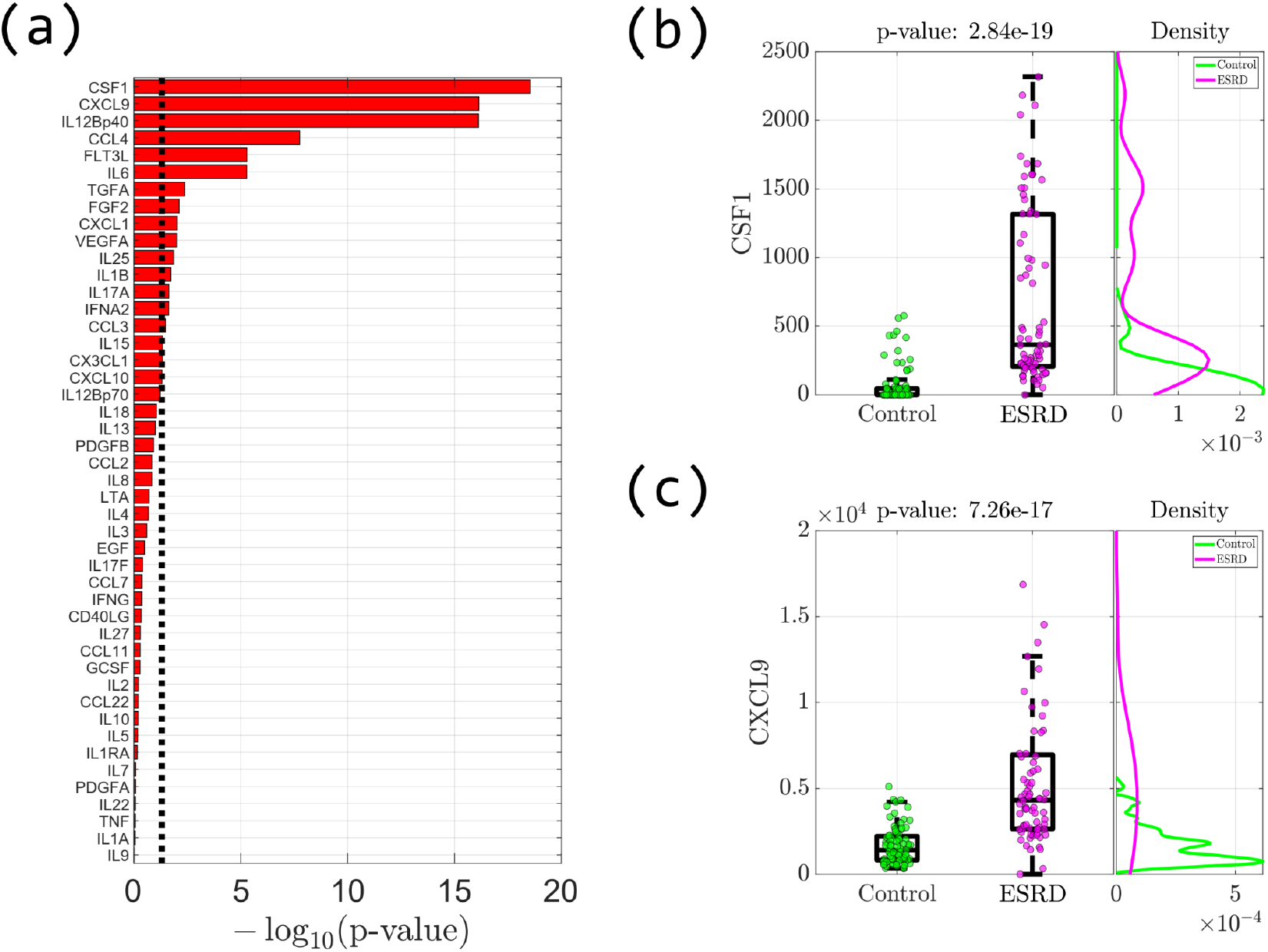
ESRD-associated inflammatory/immunological proteins. (a) Mann–Whitney U test p-value, which identifies a significant difference of biomarkers values between groups. The dashed line corresponds to 0.05. (b), (c) are two biomarkers with the highest association with ESRD: CSF1 and CXCL9.

We also tested the individual proteins for their association with chronological age, epigenetic age, phenotypic age and ipAGE in the control and ESRD groups separately (Supplementary Table 8, Figure 7(a)). Notably, some biomarkers implemented in Immunological clocks do not have a significant correlation with age by themselves.

**Figure 7.**
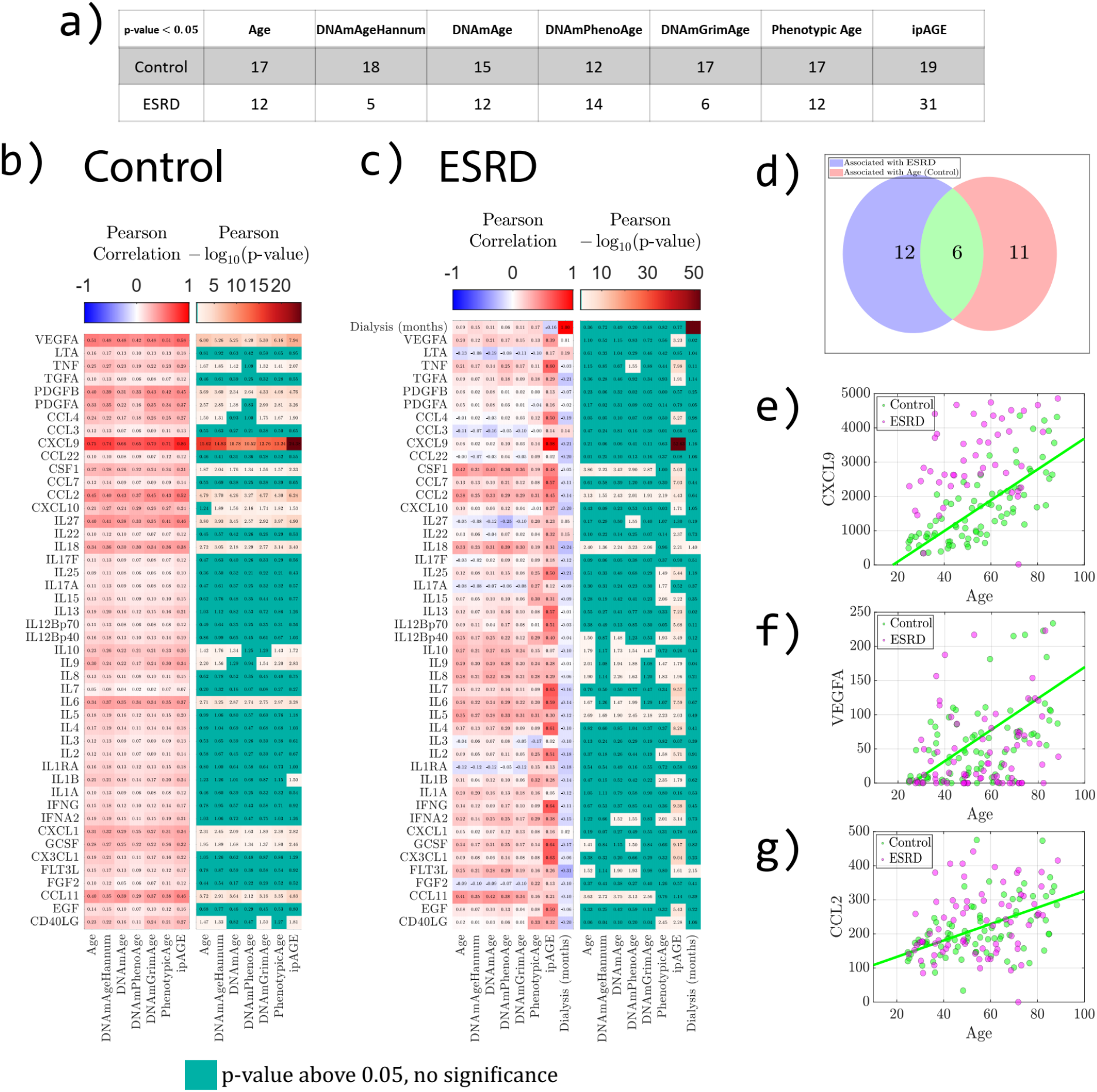
Association of inflammatory/immunological proteins with age. (a) Number of proteins significantly associated with chronological age, epigenetic ages, phenotypic age and ipAGE. (b) Pearson correlation coefficients and corresponding p-values resulting from the correlation of protein levels and each type of age, in the control and (c) ESRD groups. (d) Venn diagram of ESRD-associated and chronological age-associated inflammatory/immunological proteins in the control group. (e-g) Examples of biomarkers associated with chronological age.

The heatmaps of Pearson correlation coefficients, as well as corresponding p-values for the considered biomarkers are shown on Figure 7(b) for the group of controls and on Figure 7(c) for the patients with ESRD.

There are eleven common biomarkers associated with chronological age, epigenetic clocks, phenotypic age, and ipAGE in the control group (Figure 7(b), Supplementary Table 9). In the ESRD group only CCL2 and IL5 have significant associations with chronological, epigenetic ages, phenotypic age, and ipAGE (Figure 7(c)).

The time on dialysis, measured in months, appears as one of the variables in the ESRD group (Figure 7(c): upper row and right column). No correlation was found between time on dialysis and chronological age, epigenetic clocks, phenotypic age, and ipAGE. Only IL18 and FLT3L have significant associations with time on dialysis.

Figure 7(d) illustrates the Venn diagram for disease-associated biomarkers and chronological age-associated biomarkers in the control group. Six cytokines lie in the intersection (CCL4, CSF1, CXCL1, CXCL9, IL6, VEGFA). The most representative biomarkers associated with chronological age are shown on Figure 7(e-g).

We also considered associations between biomarkers and different types of age acceleration (Supplementary Figure 3, Supplementary Table 8). In the control group as well as in ESRD group, there are no biomarkers that are correlated with all types of age accelerations (Supplementary Figure 3(b,c)). However, in the ESRD group, most biomarkers are significantly associated with ipAGE acceleration (Supplementary Figure 3(c)). There are five biomarkers (CCL4, CSF1, CXCL1, CXCL9, IL6, VEGFA) that are associated with the disease, chronological age in the control group, and ipAGE acceleration in ESRD group (Supplementary Figure 3(d)).

Supplementary Figures 4, 5, 6 illustrate associations of all biomarkers with chronological age, ipAGE and ipAGE acceleration correspondingly. Associations between all immunological biomarkers in Control and ESRD groups are separately shown in Supplementary Figure 7.

## Discussion

In this study, we evaluated biomarkers of age in patients with end-stage renal disease, focusing on epigenetic and inflammatory/immunological biomarkers.

So far, only few studies have evaluated epigenetic changes related to kidney function and CDK^47–50^. More recently, Matías-García et al. investigated the association between epigenetic clocks and renal function^29^, reporting that epigenetic age acceleration was associated with low renal function. The cohort assessed by Matías-García et al. did not include patients with ESRD or under dialysis, that on the contrary are the focus of our study. We found that most of the epigenetic clocks, in particular DNAmPhenoAge and GrimAge, detected age acceleration in ESRD group, thus confirming and extending the results by Matías-García et al in a more extreme phenotype. We also found that EEAA is not significantly different between ESRD and controls, suggesting that immunosenescence is not a main trigger of the accelerated aging in the patients group.

Phenotypic age revealed an even more marked age acceleration in ESRD patients, with most of its components showing significantly different values compared to the control group. This indicates that ESRD produces a significant impact on a variety of homeostasis regulation processes in the body, and its effect is not limited to an increase in creatinine, the renal filtration efficiency marker. Noteworthy, while chronic inflammation is viewed as one of the pathological mechanisms of CKD and ESRD development, the ESRD group did not show an increase in C-reactive protein, the main marker of inflammation in Phenotypic Age, and lymphocyte counts in patients with ESRD were statistically lower than the control values (p=1.23e-06), suggesting the existence of more complex and systemic changes.

The development of CKD may be associated with chronic inflammatory reactions of an infectious nature (glomerulonephritis – 30% of cases) or non-infectious processes (type 2 diabetes mellitus – 55% of cases). Quite rarely, it is the result of genetically determined processes or toxic damage (15%). However, the process of CKD development is always associated with the so-called uremic inflammation. Uremic inflammation is related to the altering effects of the uremic milieu on the immune systems^51–53^, and includes mechanisms of both immunoactivation and immunosuppression. As a rule, immunoactivation during the development of uremia is associated with an increase in the synthesis of IL-1, IL-6 and TNF^51,54^. In ESRD, the innate immune system, that involves monocytes, macrophages, granulocytes, and also endothelial cell activation, is activated together with a depletion of natural regulatory T-cells, resulting in systemic inflammation and an enhanced oxidative stress. These processes are connected with an adaptive immune deficiency due to the reduction of naïve and central memory T-cells and B-cells, dendritic cells, and altered functions of polymorphonuclear leukocytes and monocytes^55^. ESRD is characterized by systemic persistent inflammation that does not subside even after complete renal sclerosis. Inflammation becomes maladaptive, uncontrolled and persistent^52^. This process recalls what is observed during inflammaging, the chronic, sterile, low-grade inflammation that develops during aging and contributes to the pathogenesis of age-diseases related^3^. Therefore, it calls for considering markers associated with inflammation, and constructing novel (immunological) clocks on their basis, that would potentially be more sensitive to disease in terms of manifested age acceleration.

We therefore developed an inflammatory/immunological clock that estimates chronological age in healthy subjects and assesses the contribution of inflammaging to the processes of age acceleration. Implementation of machine learning algorithms allowed to select meaningful immunological biomarkers and to derive an ipAGE clock model, that takes into account the complex relationships and redundancy of the cytokine network. The clock MAE is about 6 years, competing with epigenetic clocks in terms of accuracy. The components of the ipAGE clock largely overlap with the iAge clock, recently developed by Sayed et al using alternative experimental and analytical approaches^7^. While the iAge predictor was shown to be associated with cardiovascular aging, here we report that ipAGE successfully detects accelerated aging in ESRD patients.

Similar to the Phenotypic Age, ipAGE demonstrates a greater sensitivity in assessing adaptive potential of an organism compared to epigenetic clocks, due to the higher blood parameters’ variation in comparison with DNA methylation changes. In addition, in analogy to Phenotypic Age, a higher variance was observed in ESRD patients. Unlike progressive CKD, ESRD is characterized by a state of “dynamic intoxication”, an increase in toxic low molecular weight products between hemodialysis sessions. Chronic dynamic intoxication is undoubtedly a unique phenomenon because of the permanent changes of milieu intérieur and dramatic changes in the concentrations of all low molecular weight compounds. Aging is a heterogeneous process of the body’s adaptation to the changing conditions of the external environment, or, as in the case of CKD, the internal one, when a variety of genetically determined mechanisms are triggered, determining the body’s individual response and an increase in the variance of the body’s metabolic parameters with age. Thus, an increase in the age-related variability of biological markers is further stimulated by impaired homeostasis under “dynamic intoxication” conditions.

We did not find any correlation between the age acceleration evaluated according to the epigenetic clocks and ipAGE in either the ESRD group or the control group. This is in line with previous results showing that different biomarkers of age can be informative of different aspects of aging^56,57^, and further suggests that currently available age estimators perform well for population data but are still not quite suited to characterize individuals, confirming the necessity of referring to several clocks to improve reliability of results^12^. Interestingly, a positive correlation ipAGE with both pro-inflammatory and anti-inflammatory markers was found in ESRD group, pointing out the extreme complexity of the processes of homeostatic regulation of immune system. This explains the huge variance of indicators and ipAGE observed in the group of ESRD patients.

We also identified 18 plasma proteins that have significantly different levels between ESRD patients and healthy controls. Most of these proteins are part of the ipAGE clock, and 6 of them (CXCL9, VEGFA, IL6, CXCL1, CCL4, CSF1) also show significant age-association in healthy controls from our cohort. These findings indicate that a considerable number of immune system cytokines are both age and chronic kidney disease-associated, in accord with inflammaging theory.

A notable example is CXCL9 (MIG), which is associated with chronological, epigenetic, phenotypic age and with ipAGE, is one of the main components of the iAge clock from Sayed et al., and whose levels are significantly different between ESRD patients and healthy controls. For a long time, СХСL9 was not considered among the main markers of inflammaging. However, there is growing evidence of its key role in the development of age-related diseases, such as cardiovascular pathology^58^, neurodegeneration^59^, glaucoma^60^ and depressed anti-tumor immunity^61^. CXCL9 is one of the chemokines that plays a role in the induction of chemotaxis, promotes leukocyte differentiation and proliferation, and as a member of the IFN-γ-dependent cytokine family promotes the development of Th1-mediated pro-inflammatory responses. Despite the fact that the molecular and cellular action mechanisms of this chemokine are broadly known, its role in inflammaging requires further study.

Similarly to CXCL9, also CSF1 is known to increase its concentration during aging and a number of age-related diseases, such as Alzheimer’s disease, cataracts, and cardiovascular disease^50^. Other biomarkers (IL12Bp40, FGF2, FLT3L) emerged from our study have been associated with inflammation processes only, but not with aging.

Interestingly, only one of the classic markers of uremic inflammation, namely IL6, correlates in our study with age-related acceleration in people with ESRD. The terminal stage of CKD has a number of features since the hemodialysis procedure aimed at detoxifying and maintaining the body’s homeostasis is stressful: it causes dynamic changes in low molecular weight metabolic products, most of which are toxic to the human body.

Particularly interesting is the fact that most of the interleukins associated with ipAGE in the ESRD group are modulators of the B-cell immune response, while those correlated with ipAGE, chronological age and epigenetic ages in the control group are modulators of the Th1 cellular immune response. It is interesting to note the absence of correlations between individual immunomarkers and epigenetic or phenotypic age, in general, and a satisfactory performance of ipAGEclocks required 38 parameters. We conjecture that the observed variability of immunoreactivity is underpinned by the heterogeneity of the body’s adaptive responses.

In conclusion, we demonstrated that epigenetic clocks, phenotypic clock and a newly developed inflammatory/immune clock coherently detect an accelerated aging phenotype in ESRD patients, although they poorly correlate with each other. Future studies should integrate these different clocks, which are likely to grasp different aspects of human aging, in order to create new tools that could identify pathological deviations from aging-related trajectories. Furthermore, the possibility to include disease-specific parameters will allow to use these new tools at the bedside, helping in the clinical management of highly prevalent conditions in the elderly like CKD.

## Author contributions

IY, EK, MGB, CF, MV, MI contributed to the conception and design of the study. IY, EK, AK, MK, NL, MV organized the datasets. IY, AK, MK, MGB performed the statistical analysis. IY, EK, AK, MGB, CF, MV, MI wrote the manuscript. All authors contributed to manuscript revision, and read and approved the submitted version.

