## Supplementary figures and images for "Accelerated epigenetic aging and inflammatory/immunological profile (ipAGE) in patients with chronic kidney disease"

### Supplementary Figure 1

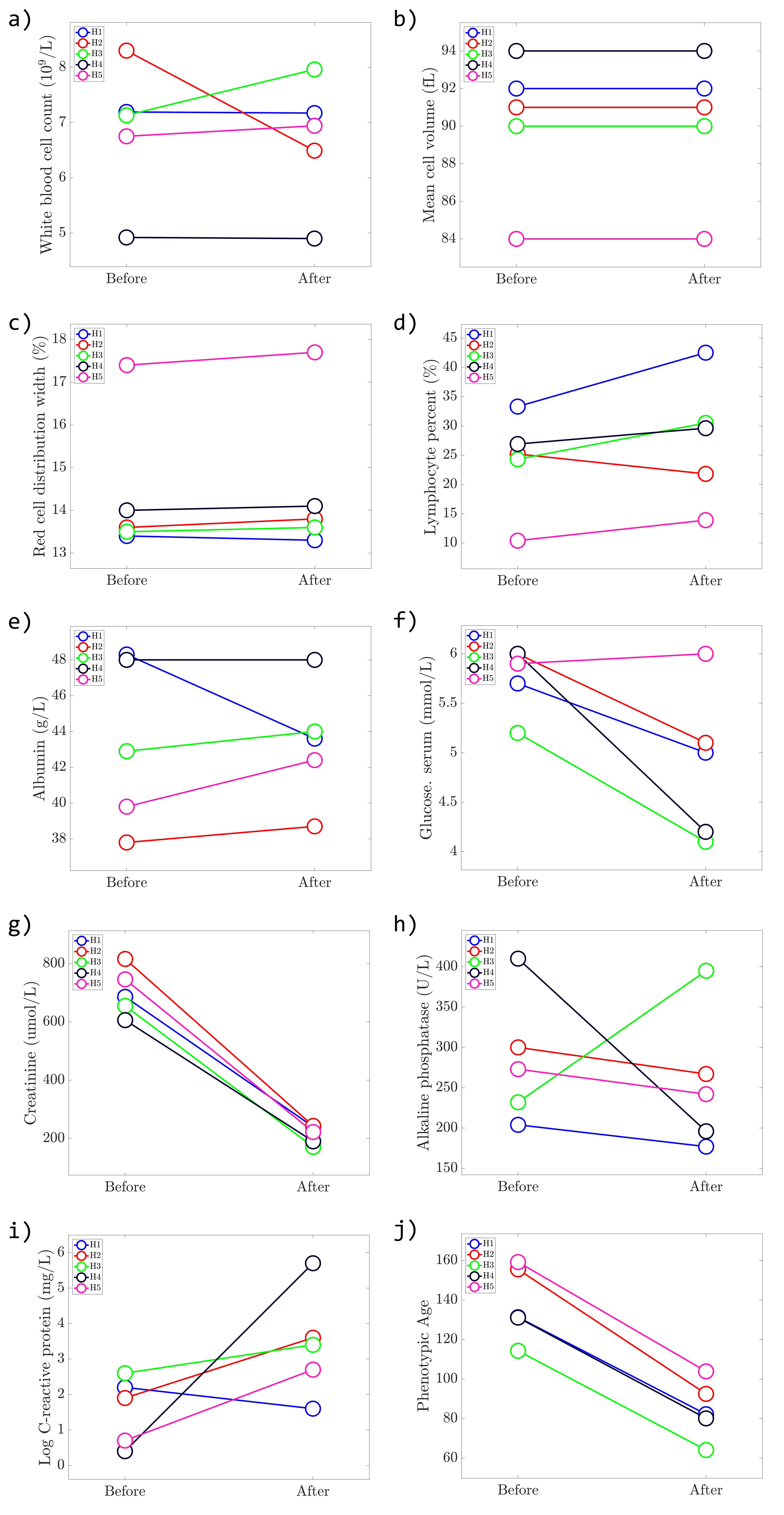

### Supplementary Figure 2

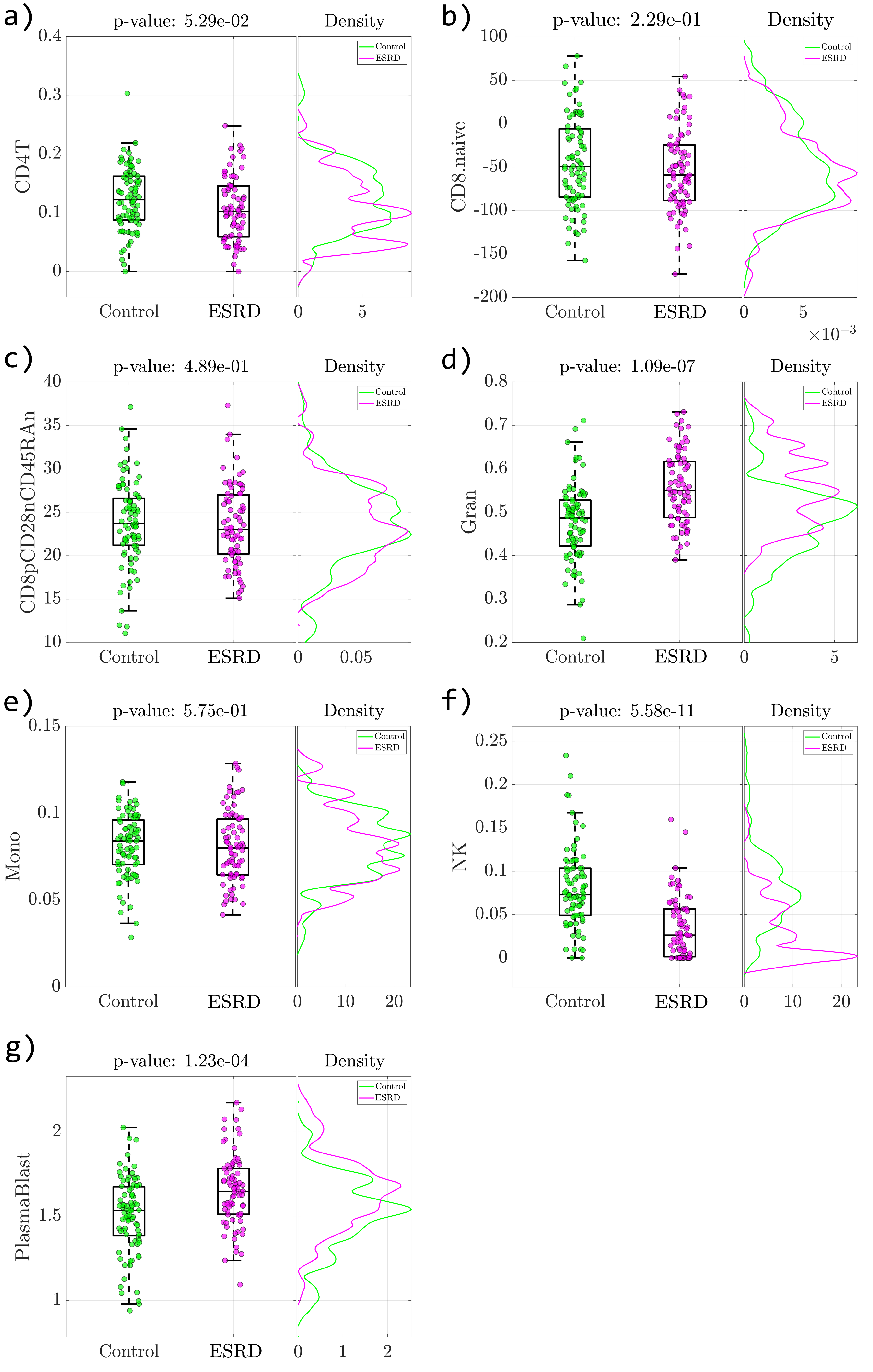

### Supplementary Figure 4

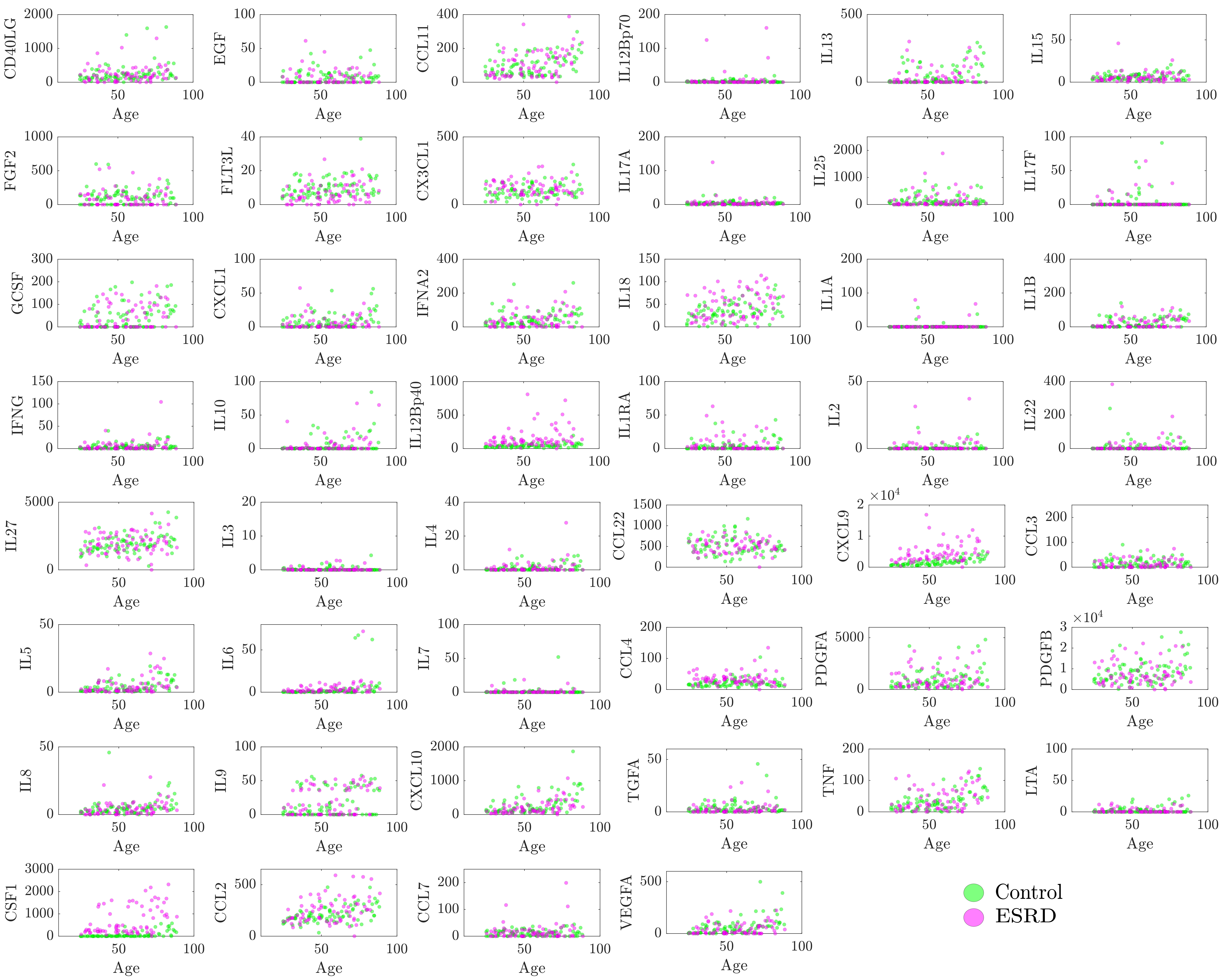

### Supplementary Figure 5

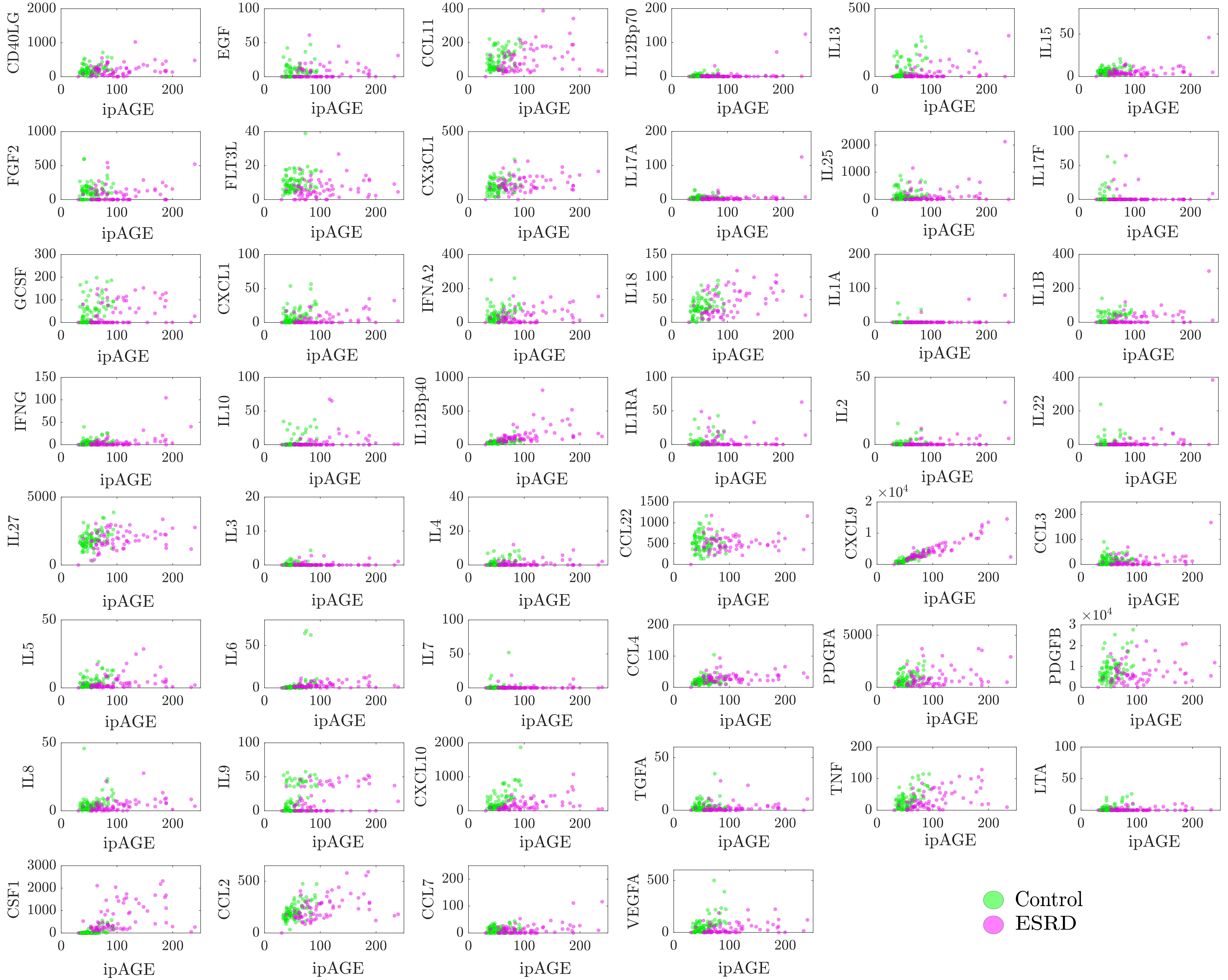

### Supplementary Figure 6

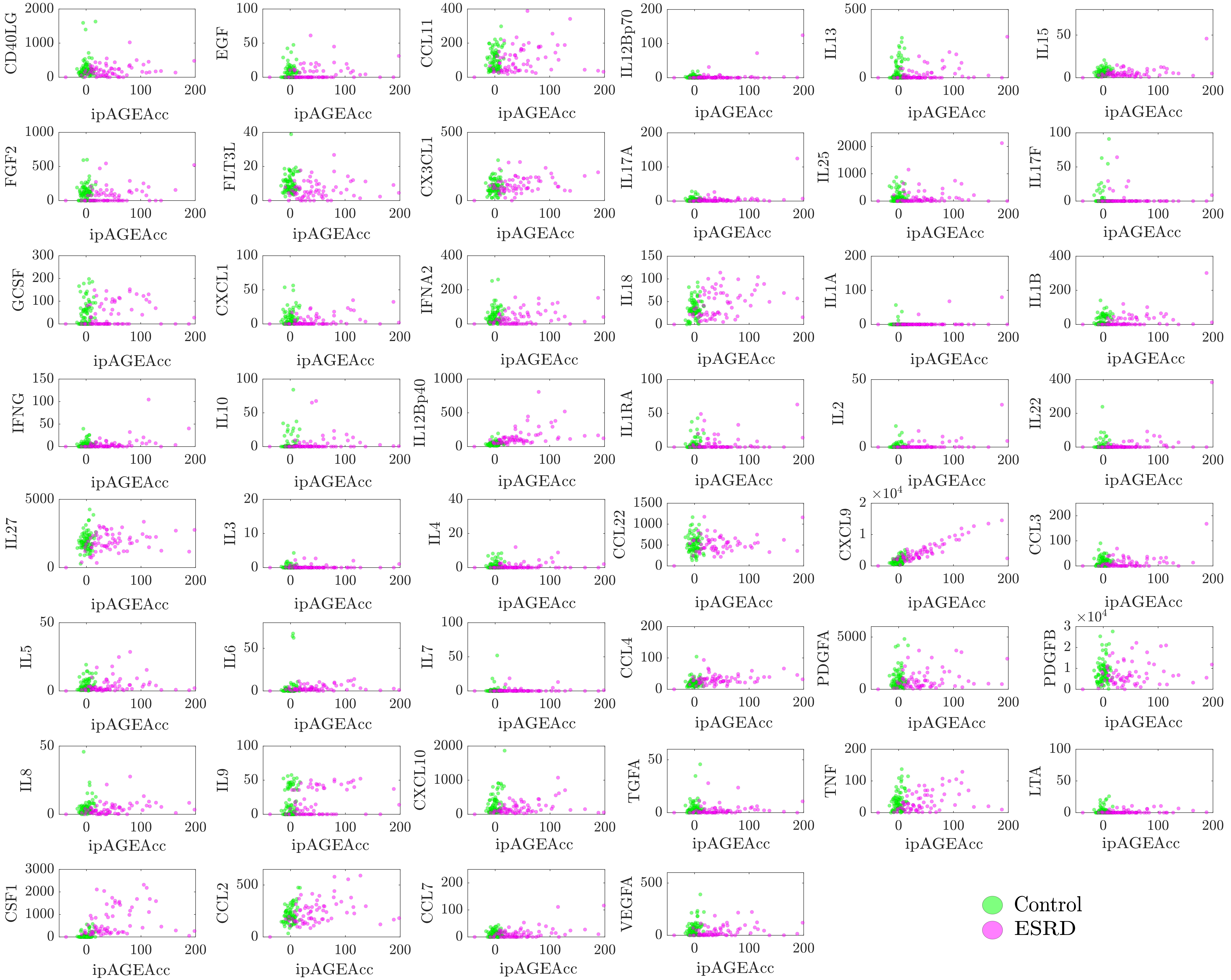

### Supplementary Figure 7

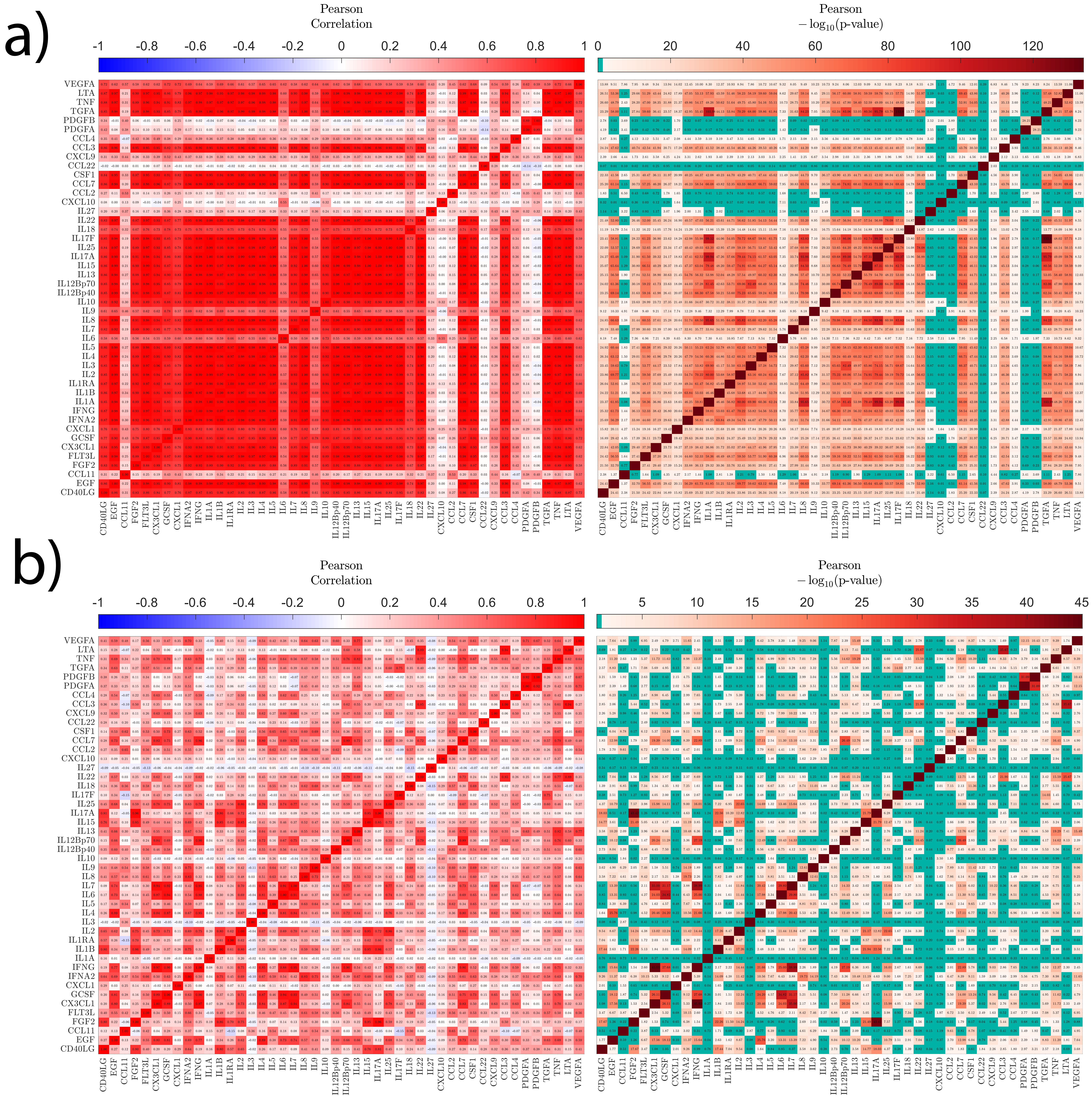
