## Supplementary Figure 3 for "Accelerated epigenetic aging and inflammatory/immunological profile (ipAGE) in patients with chronic kidney disease"

a)

| p-value < 0.05 | DNAmAgeHannum | DNAmAge | IEAA | DNAmPhenoAge | DNAmGrimAge | Phenotypic Age | ipAGE |
|---|---|---|---|---|---|---|---|
| Control | 0 | 0 | 1 | 2 | 0 | 1 | 6 |
| ESRD | 7 | 1 | 1 | 1 | 6 | 9 | 32 |

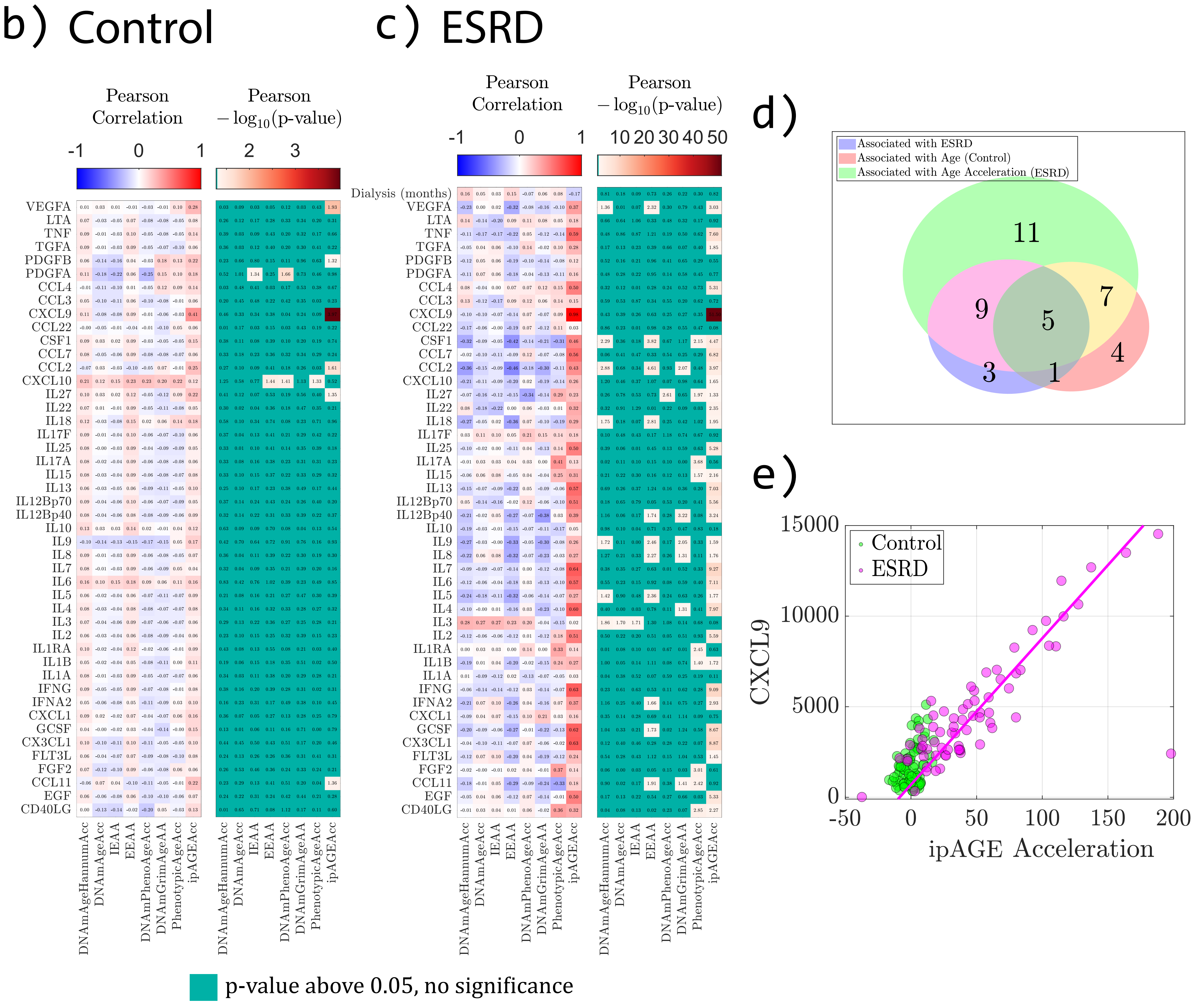
