## Supplementary Table 9 for "Accelerated epigenetic aging and inflammatory/immunological profile (ipAGE) in patients with chronic kidney disease"

Supplementary Table 9. 11 biomarkers which are associated with all types of ages

| **Biomarker** | **Name** | **Role of inflammation** | **Аge-associated** |
| --- | --- | --- | --- |
| **CXCL9** | Chemokine (C-X-C motif) ligand 9, Monokine induced by gamma interferon (MIG) | Proinflammatory chemokine → Th1 recruitment | Increases with age^1,2^  Inflammaging^3^  CVDs^4^ |
| **VEGFA** | Vascular endothelial growth factor A | Stimulation of migration of monocytes | Decreased angiogenesis^5^ |
| **CCL2** | C-C motif ligand 2, Monocyte Chemoattractant Protein 1 (MCP-1) | - The strongest factor in attracting monocytes  - Egress and migration of cells from the hematopoietic organs | Increases with age^6^  Osteoarthritis^7,8^  Alzheimer's disease^9^  CVDs^10^ |
| **IL27** | Interleukin 27 | - Аnti-inflammatory → IL10↑  - Expressed by antigen presenting cells → induces differentiation of the diverse populations of T cells | Violation of the mechanisms of anti-inflammatory action with age^11^ |
| **CCL11** | C-C motif chemokine 11, Eosinophil chemotactic protein and eotaxin-1 | Eosinophil recruitment →allergic responses | Increases with age^12^  Suppresses neurogenesis^13^  Stimulates neurodegeneration and dementia^12^ |
| **PDGFB** | Platelet-derived growth factor subunit B | No information | Neutralization of neurotoxins  Functioning of the blood-brain barrier^14^  Stimulating osteoblastoenesis^15^ |
| **IL18** | Interleukin 18, Interferon-gamma inducing factor | - Acts on CD4, CD8 T cells and NK cells to induce IFNγ production, type II interferon →Activating the macrophages  - Differentiation of naive T cells into Th2 cells  - Stimulates mast cells and basophils  - Activating apoptosis | Oncology^16,17^  Atherosclerosis^18^  Metabolic syndrome^19^  Allergy^18^ |
| **IL6** | Interleukin 6 | - Acute phase protein  - The production of neutrophils in the bone marrow  - The growth of B cells | Increases with age^20^  Atherosclerosis^21^  CVDs  Inflammaging^20,22^ |
| **CXCL1** | chemokine (C-X-C motif) ligand 1 | Chemoattractant for neutrophils | Wound healing  Tumorigenesis^23,24^ |
| **GCSF** | Granulocyte colony-stimulating factor, colony-stimulating factor 3 (CSF 3) | - Produce granulocytes  - Egress granulocytes | Age-related immunodeficiency^25^  Sacropenia^26^ |
| **CSF1** | colony stimulating factor 1, macrophage colony-stimulating factor (M-CSF) | - Proliferation, differentiation, and survival of monocytes, macrophages, and bone marrow progenitor cells  - Macrophages and monocytes→increased phagocytic and chemotactic activity | Alzheimer's disease^27^  Dementia^28,29^  Increases with age^30^  Regulate osteoclastogenesis^15^ |
